# *In vivo* characterisation of endogenous cardiovascular extracellular vesicles in larval and adult zebrafish

**DOI:** 10.1101/742692

**Authors:** Aaron Scott, Lorena Sueiro Ballesteros, Marston Bradshaw, Ann Power, James Lorriman, John Love, Danielle Paul, Andrew Herman, Costanza Emanueli, Rebecca J. Richardson

## Abstract

**Objective:** Extracellular vesicles (EVs) facilitate molecular transport across extracellular space, allowing local and systemic signalling during homeostasis and in disease. Extensive studies have described functional roles for EV populations, including during cardiovascular disease, but the *in vivo* characterisation of endogenously produced EVs is still in its infancy. Due to their genetic tractability and opportunities for live imaging, zebrafish represent an ideal but under-used model to investigate endogenous EVs. The overall aim of this study was to establish a transgenic zebrafish model to allow the *in vivo* identification, tracking and extraction of endogenous EVs produced by different cell types.

**Approach and Results:** Using a membrane-tethered fluorophore reporter system, we show that EVs can be fluorescently labelled in larval and adult zebrafish and demonstrate that multiple cell types including endothelial cells and cardiomyocytes actively produce EVs *in vivo*. Cell-type specific EVs can be tracked by high spatiotemporal resolution light-sheet live imaging and modified flow cytometry methods allow these EVs to be further evaluated. Importantly, we demonstrate the utility of this model by showing that cardiomyocytes, endothelial cells and macrophages exchange EVs in the adult heart and that ischaemic injury models dynamically alter EV production.

**Conclusions:** We have developed a powerful *in vivo* zebrafish model for the investigation of endogenous EVs in all aspects of cardiovascular biology and pathology. A cell membrane fluorophore labelling approach allows cell type-specific tracing of EV origin without bias towards the expression of individual protein markers and will allow detailed future examination of their function.

## Introduction

Extracellular vesicles (EVs) are plasma membrane-bound particles produced and released by most cell types. EVs can be split into three classes: exosomes, formed from an endocytic pathway, microvesicles that are shed from the cell surface, and apoptotic bodies, which are shed from cells undergoing apoptosis ^1, 2^. EVs can be trafficked locally and systemically and have been isolated from various biological fluids, including blood ^3, 4^ and pericardial fluid ^5, 6^. Various surface glycans, lipids and proteins reportedly guide EVs to regions of extracellular matrix or recipient cells, and these interactions play integral roles in communicatory pathways ^7^. EVs play roles in homeostasis ^8, 9^ and are implicated in the progression of many diseases, including cardiovascular disease ^10, 11^. EVs have been reported to facilitate communication between multiple cell types within the cardiac microenvironment, including cardiomyocytes, fibroblasts, immune and endothelial cells (ECs) ^12–15^ and some populations of EVs are thought to be proangiogenic and cardioprotective ^11, 15, 16^.

Zebrafish have emerged as a powerful model for cell/developmental biology and human diseases and have many advantages including high fecundity, rapid external development, genetic tractability and unrivalled cellular level *in vivo* imaging ^17^. Most zebrafish studies have been performed at larval stages, however, the advantages offered by adult zebrafish, including retained regenerative capacity, have led to an increasing number of studies using this model ^17^. Clinically relevant models of tissue injury that allow subsequent evaluations of regenerative processes are well established in adult zebrafish ^18^. In the heart, correct cardiac regeneration, including complete cardiomyocyte renewal, has been shown to be reliant on multiple different cell types including inflammatory cells and ECs ^19–22^, implying the need for a complex communication system. Little is known about the role of endogenous EVs in regenerative contexts despite the potential to identify pro-regenerative signals being exchanged between cell types.

The potential of EVs as biomarkers of disease and as novel therapeutic delivery vehicles has generated significant interest in recent years ^4, 23^. However, the ability to reliably define the heterogeneous spectrum of endogenous EV subtypes and their functional significance *in vivo* is still in its infancy ^2^. The majority of EV characterisation to date, by necessity, has been performed *in vitro* ^24^. Recent studies have developed novel ways to label exogenous EVs allowing their biodistribution and functional roles to be investigated *in vitro* and *in vivo* ^5, 25–27^. Whereas these studies using exogenous EVs are beginning to identify important roles in multiple tissues and disease states, they are unable to address the full complexity of endogenous *in vivo* EV populations. Recent reports have started to bridge this gap by investigating endogenously produced EVs in larval zebrafish, including apoptotic bodies ^28^ and CD63+ exosomes ^29^. However, *in vivo* studies of endogenous EVs released by ECs and cardiomyocytes under homeostatic and disease conditions have not been attempted so far.

Here, we describe techniques using stable transgenesis to fluorescently label native cell-type specific EVs *in vivo* in larval and adult zebrafish, allowing these vesicles to be tracked, extracted and validated. We demonstrate that global zebrafish EVs as well as EC- and cardiomyocyte-derived EVs can be observed in the peripheral circulation and the pericardial space, respectively. We report that using adapted flow cytometry techniques these cell-type specific EVs can be analysed and purified from tissue samples and evaluated *ex vivo*. Importantly, we describe the transfer of EVs between different cell types resident in the adult heart. Finally, we have used models of tissue injury in larval and adult zebrafish to demonstrate dynamic changes to the EV profile.

## Materials and Methods

Please also see the Major Resources Table in the Supplemental Materials.

### Zebrafish lines and procedures

The Tg(*actb2:HRAS-EGFP*) ^30^, Tg(*tbp:GAL4*);(*UAS:secA5-YFP*) ^31^The Tg(*actb2:HRAS-EGFP*) ^30^, Tg(*tbp:GAL4*);(*UAS:secA5-YFP*) ^31^, Tg(*kdrl:mCherry-CAAX*) ^32^, Tg(*myl7:GFP*) ^33^, Tg(*fli1:EGFP*) ^34^, Tg(*mpeg1:EGFP*)gl22 ^35^ and Tg(*myl7:HRAS-mCherry*) ^32, 36^, Tg(*myl7:GFP*) ^33^, Tg(*fli1:EGFP*) ^34^, Tg(*mpeg1:EGFP*)gl22 ^35^ and Tg(*myl7:HRAS-mCherry*)^36^ lines have been described previously. In all cases, animals were anaesthetised via immersion in 0.025% MS-222 (Sigma; A5040) and euthanised via immersion in an overdose of anaesthetic. For hypoxia experiments, 3 days post fertilisation (dpf) larvae were incubated at 28°C in Danieau’s buffer in a hypoxia workstation (InvivO_2_ 300, Ruskinn) in a 5% CO_2_/95% N_2_ gas mixture to maintain 5% oxygen ^37^ for 18 hours. Cardiac cryoinjuries on adult zebrafish were carried out as described previously ^38^. Briefly, fish were anaesthetised and placed ventral side up in a pre-cut sponge soaked in aquarium water containing anaesthetic. A 4 mm incision was made directly above the heart to expose the ventricle, which was dried using a sterile cotton swab and a liquid nitrogen cooled probe was applied for 30 seconds. Adult fish used were aged 4 - 18 months and were randomly assigned to experimental groups. All lines were maintained according to standard procedures and all animal work was carried out in accordance with UK Home Office and local University of Bristol regulations.

### Imaging

Larval and adult zebrafish were anaesthetised and mounted in 1% low-gelling agarose (Sigma). Live imaging was performed on a ZEISS Lightsheet Z.1 system with a 40x W Plan Apochromat objective or a Leica TCS SP8 AOBS confocal laser scanning microscope with a 25x/0,95 W HC FLUOTAR objective with resonant scanner (frame intervals of 0.02-0.04 seconds). To image free particle movement in larvae, synthetic EVs containing a Cy5 conjugated microRNA (cel-miR-39-3p; not present in zebrafish ^39^) were microinjected into the pericardial space. Adult hearts were dissected, fixed in 4% Paraformaldehyde and embedded and imaged as above with the conventional confocal scanner. Images were processed using Fiji ^40^, IMARIS (version 9.5.0, Oxford Instruments, United Kingdom) and Huygens Professional (version 16.10, Scientific Volume Imaging, The Netherlands). Deconvolved images are noted in the figure legends. For manual counts of EVs, all analysis was blinded and positive events were counted from 1-minute videos of the peripheral circulation.

### Cell Dissociation and EV Isolation

Adult hearts (1-3 hearts per experiment) or 4-5 dpf whole larvae (16-20 per experiment) were dissociated in perfusion buffer (PBS plus 10 mM HEPES, 30 mM Taurine and 5.5 mM Glucose) plus 0.25% Trypsin, 12.5 μM CaCl_2_ and 5 mg/ml Collagenase II (Worthington Biochemical Corp; LS004176) for 1 hour at 32 °C. Reactions were stopped with perfusion buffer plus 10% (vol/vol) FBS and 12.5 μM CaCl_2_. Dissociated cells were centrifuged at 300 *g* (2 x 10 minutes), 1200 *g* (2 x 10 minutes) and passed through a 0.8 μm sterile filter. Crude EVs were obtained by centrifugation at 20,000 *g* (30 minutes) or 100,000 *g* (70 minutes) (Optima XPN Ultracentrifuge, SW 32 Ti Rotor, Beckman Coulter) for large and/or small populations, respectively, and resuspended in 300 μl sterile filtered PBS. Samples were either used immediately or snap frozen in liquid nitrogen, stored at −80°C and later thawed at 37°C for 2 minutes before use.

### Dynamic Light Scatter (DLS) and Nanoparticle Tracking Analysis (NTA)

Hydrodynamic particle size of crude EV samples in PBS was measured in triplicate using a Zetasizer Nano-ZS (Malvern Instruments, Malvern Hills, UK). Particle concentration and size distribution were determined using a ZetaView NTA system (Particle Metrix, Germany) and ZetaView software (version 8.05.11 SP1). 100 nm polystyrene standard particles were used for calibration measurements. For video acquisition: sensitivity = 85, shutter speed = 100, acquisition = 30 frames per second. Each sample was measured at 11 different positions, with 2 cycles of readings at each position. After automated analysis and removal of any outliers from the 11 positions, the size (diameter in nm) and the concentration (particles/mL) were calculated.

### Flow Cytometry

From crude EV samples, intact EVs were labelled with calcein violet 450 AM (eBioscience; 65-0854-39) as previously described ^41^. Detergent treated samples had 0.05% Triton X-100 (Sigma; T8787) added. Flow cytometry analysis and fluorescence activated vesicle sorting (FAVS) was performed on a BD Influx system, with optimisations to allow for the reliable detection of EVs ^42^. Briefly, 200 mW 488 nm (small particles and GFP), 50 mW 405 nm (calcein 450) and 100 mW 552 nm (mCherry) lasers were used with bandpass filters: 530/40 nm, 460/50 nm and 610/20 nm. A small-particle detector provided high sensitivity in detecting forward scatter and a 0.45 threshold on a logarithmic scale was used. A 4 mm obscuration bar optimally detected submicron particles. For flow cytometry of EVs, a 100 μm nozzle and 21 PSI was used. To increase speed and throughput for FAVS, a 70 μm nozzle and 42.9 PSI was used. EVs were sorted into 100 μl PBS. All flow cytometry experiments were performed at 4°C.

Imaging flow cytometry analysis was carried out as previously described ^43^. Briefly, a fully calibrated (ASSIST tool) ImageStream^®x^Mk II (Amnis-Luminex, Seattle, USA) with 405/488 nm excitation lasers, a 785 nm side scatter laser, brightfield illumination and a six channel charge-coupled device camera with time delay integration was used. For maximum resolution and sensitivity, fluidics were set at low speed, magnification at 60x (0.3 mm^2^/pixel) and 1.5 μm diameter, carboxylated polystyrene microspheres (Speed beads) were run continuously during data acquisition. Data analysis was performed using ImageStream Data Exploration and Analysis Software (IDEAS^®^6.2 EMD Millipore, Seattle).

### Transmission Electron Microscopy (TEM)

Samples were prepared for TEM by established methods ^44^. Five microlitres of isolated EV sample was placed onto Formvar-carbon coated, glow-discharged, 300 mesh copper grids (prepared in house by Wolfson Bioimaging Facility staff), left for 30 seconds and the buffer removed with blotting paper. Grids were negatively stained in 1% uranyl acetate for 30 seconds, air-dried and visualised on a FEI Tecnai 12 120kV BioTwin Spirit TEM and images acquired using a FEI Eagle 4k x4k CCD camera.

### Total Internal Reflection Fluorescence (TIRF) Microscopy

EV prep for TIRF microscopy was performed as previously described ^45^. Briefly, 50 μl sorted EV samples plus 0.06 μm carboxylate-modified microspheres (Sigma; T8870) (1:10000 final concentration) were placed onto 35 mm glass bottom dishes (Mattek; P35G-1.5-14-C) and allowed to settle. Images were acquired using a Leica AM TIRF multi-colour system attached to a Leica DMI 6000 inverted epifluorescence microscope.

### Western Blot

Western blot analysis was carried out using standard methods. Briefly, cells, large EV and small EV pellets were lysed on ice for 10 min with lysis buffer (125 mM NaCl, 20 mM TRIS pH 7.4, 1% Nonidet P40 (Sigma Aldrich; 11754599001), 10% Glycerol, 0.1 mg/ml Phenylmethylsulfonyl fluoride, protease cocktail inhibitor (Sigma Aldrich; 04693159001), 50 mM NaF and 10 mM Na_3_O_4_V). Equal protein concentrations were resolved via 8% SDS–PAGE and transferred to Immobilon-P polyvinylidene difluoride membrane (Sigma Aldrich; IPVH00010). Antibodies used: anti-ALIX (Sigma Aldrich; SAB4200476, 1:500). LI-COR IRDye 800CW secondary antibody (LI-COR Biosciences; 926-32211) (1:5000), detected using an Odyssey Fc imaging system from LI-COR Biosciences (Cambridge, UK).

### Statistics

In all cases n numbers refer to biological replicates unless otherwise stated. All experiments were repeated at least twice. GraphPad Prism6/7 was used for raw data recording/analysis. Statistical tests used were nonparametric Mann-Whitney or Kruskal-Wallis/Dunn’s multiple comparison tests or a randomised permutation test using total variation distance between two groups (details in figure legends). All statistical analysis was blinded. In all cases error bars represent SD. For all data sets a Grubb’s outlier test was performed and any significant outliers (alpha = 0.05) were removed. p values are included in all graphs.

## Results

### Stable labelling and imaging of endogenous EVs *in vivo*

Despite recent advances in labelling methods, the distribution and function of cell type specific EVs *in vivo* remains incompletely understood. To label large proportions of EVs produced by individual cell types and avoid targeting sub-populations via EV enriched protein labelling, we hypothesised that stable transgenic zebrafish lines expressing cell membrane-tethered fluorophores would also label EVs produced by those cells (Figure IA in the Data Supplement). Similar unbiased inner leaflet cell-membrane labelling techniques have been used to label cells and their derived EVs in other models ^46^. To demonstrate that endogenous EVs could be labelled with this system, we initially used a transgenic line where prenylated GFP is driven by a near-ubiquitous promoter, tethering the fluorophore to the inner leaflet of the plasma membrane (Tg(*Ola.Actb:Hsa.HRAS-EGFP*) (referred to as Tg(*actb2:HRAS-EGFP*)); Figure IA,B in the Data Supplement ^30^). Live imaging of Tg(*actb2:HRAS-EGFP*) larvae revealed sub-cellular GFP+ particles in the peripheral circulation and in the pericardial space at 3 dpf (Figure IC,G and Videos I & II in the Data Supplement). The pericardial space is a relatively large extracellular space in larval zebrafish (Figure ID-F in the Data Supplement) and GFP+ EVs were observed moving within the pericardial fluid, influenced by the movement of the heart (Video II in the Data Supplement). Additionally, we used a ubiquitously expressed secreted Annexin-V line which binds with high affinity to the phosphatidylserine expressed on the outer surface of apoptotic cells (referred to as Tg(*tbp:GAL4*);(*UAS:secA5-YFP*) ^31^; Figure IIA,B in the Data Supplement), but also on populations of EVs, including those relevant in cardiovascular disease ^47, 48^. As with the *actb2* line we observed Annexin-V labelled particles in both the peripheral circulation and pericardial space of larval zebrafish (Figure IIC,D and Video III in the Data Supplement).

Although these transgenic lines suggested we could successfully identify endogenous EVs *in vivo*, they do not allow us to determine the cellular origin of these subcellular particles. We therefore made use of two more transgenic lines that express a prenylated fluorophore driven by cell specific, cardiovascular relevant promoters, whereby the fluorophore is incorporated into the EVs in a cell-type specific manner (Figures 1,2). Live confocal or light-sheet imaging of larval zebrafish expressing an endothelial specific promoter driving membrane tethered mCherry (Tg(*kdrl:mCherry-CAAX*) ^32^; revealed the presence of fluorophore-labelled subcellular particles (referred to as EC-EVs) in the peripheral circulation (Figure 1A-D and Video IV in the Data Supplement). Putative EC-EVs moved rapidly in the peripheral circulation with the blood flow and were observed both in arterial flow in the dorsal aorta (DA) (mean = 63 ± 23 (SD) EVs/minute) and in venous flow in the caudal haematopoietic tissue (CHT) (mean = 333 ± 33 (SD) EVs/minute) (Figure 1B-D and Video IV in the Data Supplement). Dual imaging with brightfield confirmed that these EC-EVs moved independently from cells in the blood (Figure 1B,C and Video IV in the Data Supplement). It has been suggested that macrophages of the innate immune system can receive EVs from multiple cell types ^25, 29^ and as macrophages are present in the peripheral circulation of larval zebrafish we investigated if they may receive EC-EVs in our model. Live imaging of Tg(*kdrl:mCherry-CAAX*); Tg(*mpeg1:EGFP*) double transgenic fish (GFP labelling macrophage cytoplasm) reveals transfer of EC-EVs to intra-vascular macrophages which were observed making protrusions into the lumen of vessels, potentially to capture passing EVs (Figure 1D and Video V in the Data Supplement), as previously shown for other EV populations ^25, 29^.

**Figure 1.**
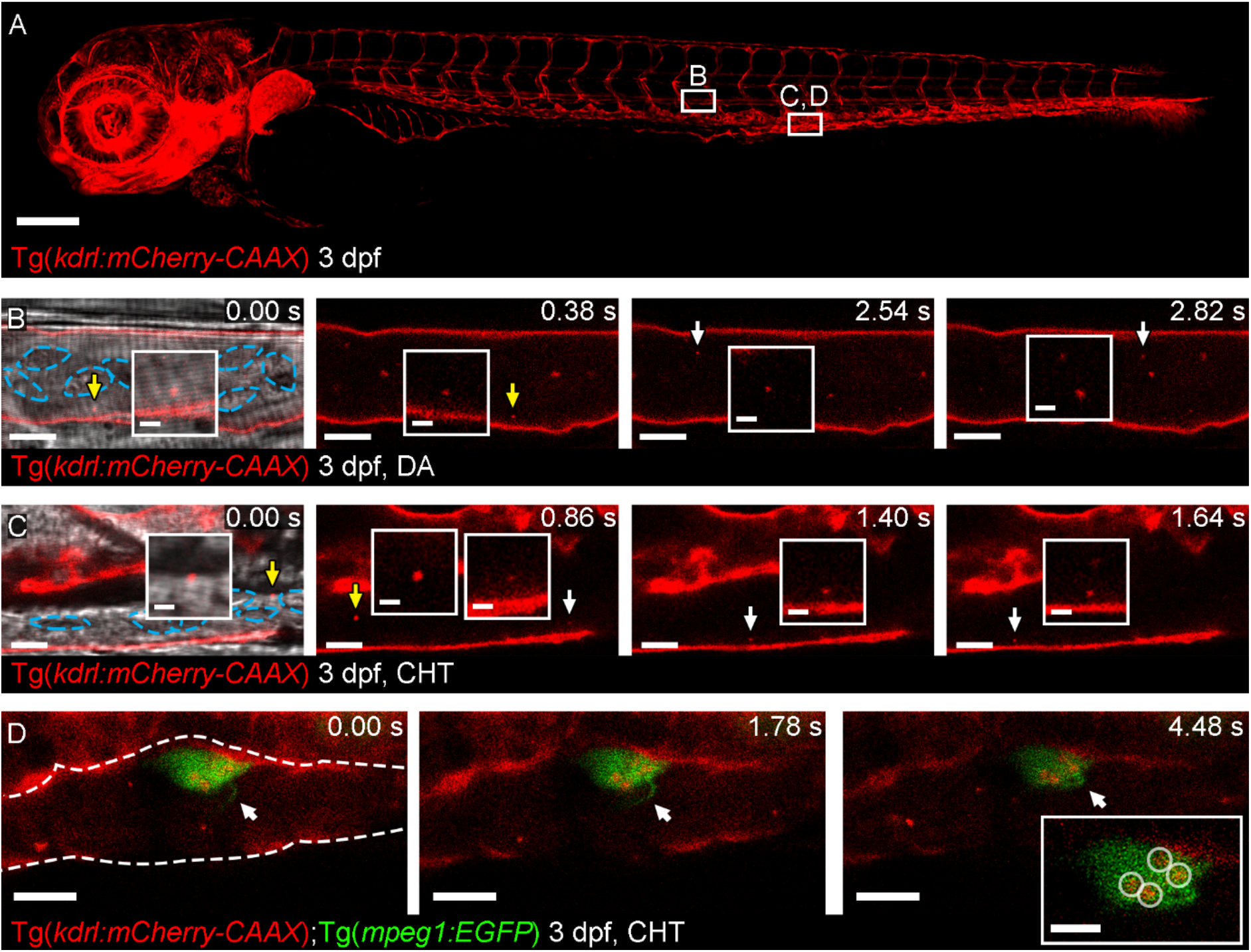
Live imaging of EC-EVs in the peripheral circulation. (**A**) Overview image of a Tg(*kdrl:mCherry-CAAX*) fish at 3 dpf. All ECs are labelled with mCherry. The boxed areas define the regions shown in the image sequences in B,C,D as labelled. (**B**) Image sequence of mCherry+ EC-EVs moving through the DA (arrows and inset). (**C**) Image sequence of mCherry+ EC-EVs moving through the CHT (arrows and inset). (**D**) Image sequence of a macrophage in the CHT of a Tg(*kdrl:mCherry-CAAX*); Tg(*mpeg1:EGFP*) fish at 3 dpf. The macrophage makes a protrusion into the vessel lumen (arrows) and intracellular compartments contain mCherry+ EC-EVs (circled in inset). The blue dashed lines in B,C demark blood cells in the brightfield image. The white dashed line in D demarks the endothelium. Anterior is to the left. Scale bars: A = 200 μm; B,C = 5 μm; insets in B,C = 2 μm; D = 10 μm; inset in D = 5 μm.

**Figure 2.**
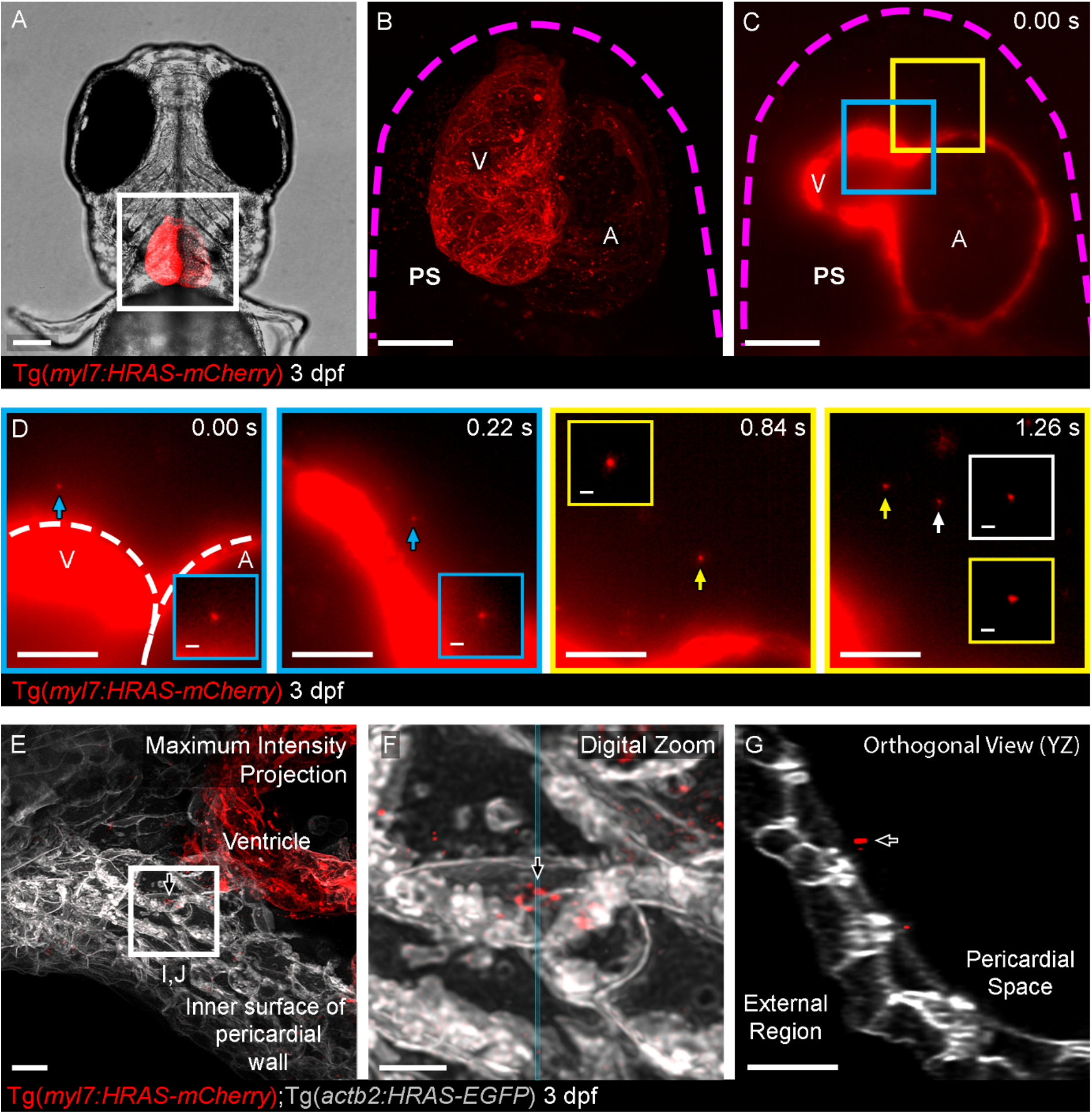
Live imaging of CM-EVs in the pericardial space. (**A**) Ventral view of 3 dpf Tg(*myl7:HRAS-mCherry*) zebrafish, the boxed region highlights the position of B,C. (**B,C**) Overview images of the entire heart of Tg(*myl7:HRAS-mCherry*) fish in ventral view. B shows a maximum projection of a fixed fish; C shows a single plane of live light-sheet imaging. (**D**) Image sequences of higher magnification views of the colour coded boxed areas in C. mCherry+ CM-EVs are observed moving through the pericardial space (arrowed and inset). (**E-G**) A maximum intensity projection of deconvolved images of the ventricle and internal surface of the pericardial wall (E) reveals static CM-EVs as shown with digital zoom of boxed region (F). The orthogonal view (YZ - blue line in F) of this region suggests the EVs are interacting with a layer of unmarked ECM rather than direct contact with underlying cells (G). Abbreviations: A = atrium; V = ventricle; PS = pericardial space. The magenta dashed line in B,C demarks the pericardial wall. The white dashed line in D demarks the ventricle and atrium. Arrows indicate EVs. Anterior is to the top in A-D and left in E,F. Scale bars: A = 100 μm; B,C = 50 μm; D = 20 μm; insets in D,F = 2 μm; E,G = 5 μm.

Live imaging of the heart of 3 dpf larvae with a cardiomyocyte specific promoter driving membrane tethered mCherry (Tg(*myl7:HRAS-mCherry*) ^36^) revealed mCherry+ particles (referred to as CM-EVs) moving within the pericardial fluid, as observed for Tg(*actb2:HRAS-EGFP*) labelled EVs (Figure 2D and Video VI in the Data Supplement). This movement is similar to that described for free proepicardial cells which dissociate from the dorsal pericardial wall and move, influenced by the heartbeat, within the pericardial fluid during epicardial formation ^49^. This process of free proepicardial cell movement is largely complete by 3 dpf ^49^, and our own data suggests the pericardial space is devoid of free moving cells at this stage ruling out interactions of these EVs with cells (Figure IIIA-D and Video VII in the Data Supplement). Additionally, we injected fluorescent ~50 nm synthetic nanoparticles ^39^ into the pericardial space of 3 dpf larvae and observed similar stochastic movement confirming the EVs were free moving (Figure IIIE and Video VIII in the Data Supplement). Some static CM-EVs were observed interacting with the pericardial wall, potentially with regions of extracellular matrix (Figure 2E-G).

### Validation of endogenous EVs

To characterise these endogenous EVs further we used a tissue dissociation protocol followed by differential centrifugation and filtration to separate a crude “large” EV fraction (centrifugation at 20,000 g) and a crude “small” EV fraction (centrifugation at 100,000 g) from entire zebrafish larvae (Figure 3A). Analysis of these different fractions using Dynamic Light Scattering (DLS) supported the isolation of different sized groups of particles (Figure 3B) with a mean diameter of 263.6 nm ± 24.38 nm (SD) and 128 nm ± 12.86 nm (SD), respectively, in line with previous reports ^50, 51^. To allow us to obtain and verify our endogenous fluorescently labelled EVs, we used a modified flow cytometer to analyse ubiquitous (*actb2*(GFP)+) EVs extracted from whole zebrafish larvae at 3 dpf (Figure 3C-H), using similar methods to those previously described ^42^. Isolated EVs were labelled with calcein violet 450 AM, which labels intact vesicles ^41^, prior to analysis by flow cytometry. Total populations of calcein+ and *actb2*(GFP)+ EVs were assessed from EVs extracted from pools of non-transgenic and Tg(*actb2:HRAS-EGFP*) larvae (Figure 3C-H). Control experiments of singly labelled EVs (GFP or calcein) from transgenic and non-transgenic zebrafish allowed double positive (*i.e*. cell type specific (fluorophore+) and intact (calcein+)) EV specific gates to be assigned, avoiding any background signal (Figure 3C-F). Analysis of EV fractions following detergent treatment confirmed their lipidaceous nature (Figure 3G,H). Flow cytometry analyses suggest that 29% ± 8.6% (SD) of total events are labelled with calcein, representing intact EVs (Figure 3H). Similarly, flow cytometry analysis suggests that 30% ± 6.9% (SD) of calcein+ vesicles are also *actb2*(GFP)+ (Figure 3F). We also analysed calcein+ EVs extracted from Tg(*actb2:HRAS-EGFP*) larvae using an imaging flow cytometry ImageStream^®x^Mk II system, confirming the presence of individual single or double labelled vesicles (Figure IVB,C in the Data Supplement). Imaging flow cytometry suggests 65% of total events are calcein(violet)+ (concentration as per routine 300 μl resuspension volume = 2.14 x 10^8^ EVs/ml) demonstrating the enhanced sensitivity and value of the ImageStream system ^52^. This analysis suggests 34% of calcein+ vesicles are also *actb2*(GFP)+, supporting our previous flow cytometry assessments.

**Figure 3.**
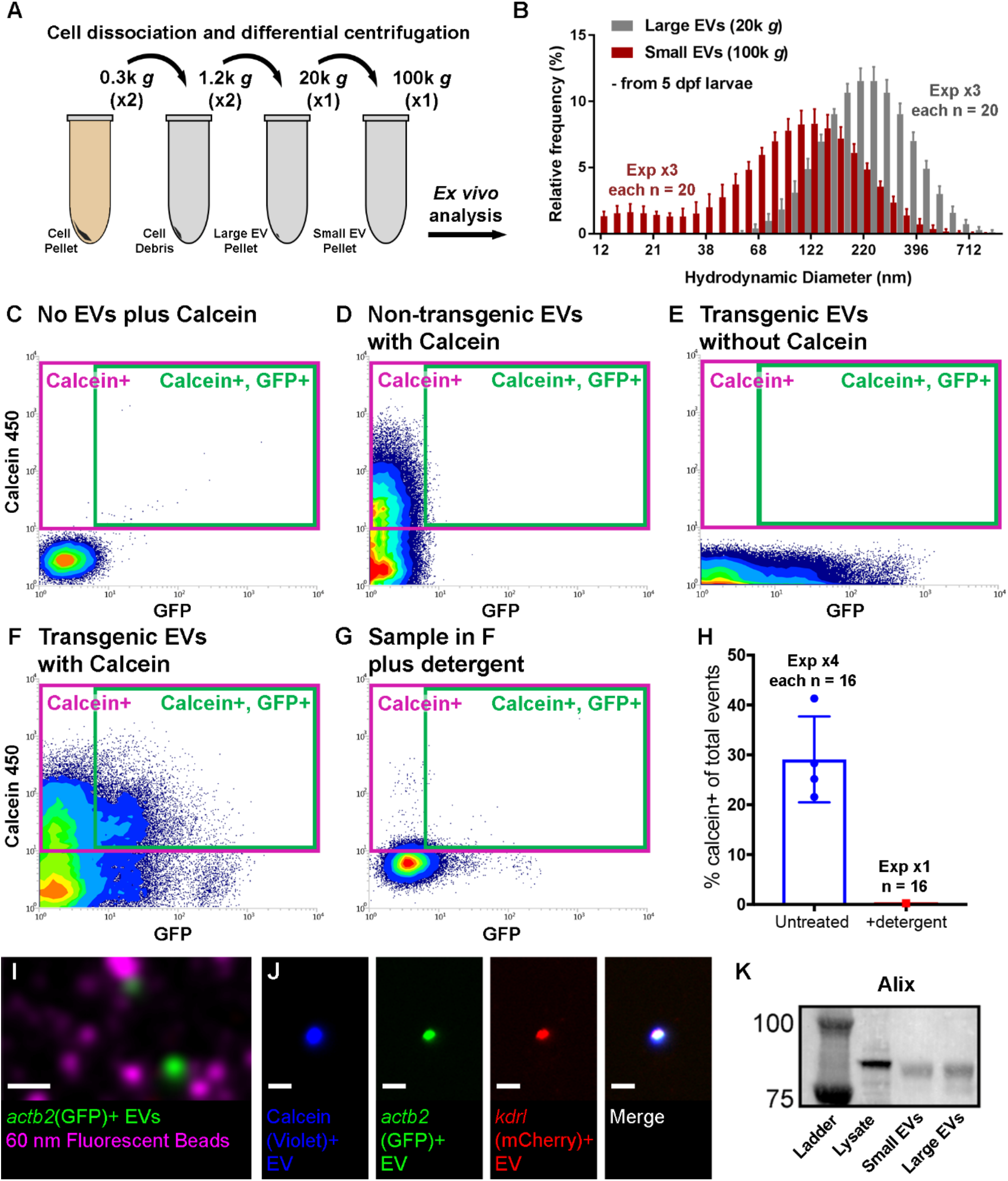
Validation of endogenous cardiovascular EVs from zebrafish using a modified FACS protocol. (**A**) Schematic describing the centrifugation steps taken to isolate EV fractions following cell dissociation of whole zebrafish larvae. (**B**) Histogram of DLS analysis of different EV fractions from whole 5 dpf larvae indicating that smaller EVs are pelleted at 100,000 *g* than at 20,000 *g*. (**C-G**) Typical flow cytometry scatter plots showing the gates used to sort EVs and the controls used to define these gates: An extraction buffer only control plus calcein AM reveals background noise (C), EVs extracted from non-transgenic (wildtype) fish and labelled with calcein AM allow us to assign a calcein+ gate (D), EVs extracted from Tg(*actb2:HRAS-EGFP*) fish without calcein AM indicates GFP+ EVs (E) and analysis of EVs extracted from Tg(*actb2:HRAS-EGFP*) fish with calcein AM labelling identifies a gate of GFP+ calcein+ EVs (F). Treatment of transgenic EVs labelled with calcein AM plus detergent destroys the majority of EVs, confirming their lipidaceous structure (G). Similar gating strategies can be used to analyse mCherry+ EVs from Tg(*kdrl:mCherry-CAAX*) and Tg(*myl7:HRAS-mCherry*) fish. (**H**) Plot of the percentage of calcein+ EVs of total events from untreated and detergent treated samples. **(I)** TIRF microscopy allows high resolution imaging of extracted EVs from Tg(*actb2:HRAS-EGFP*) fish (green) alongside 60 nm synthetic fluorescent beads (magenta). (**J**) TIRF imaging of a single EV derived from a Tg(*actb2:HRAS-EGFP*); Tg(*kdrl:mCherry-CAAX*) double transgenic fish stained with calcein AM. (**K**) Western analysis of protein extracted from cell and EV fractions confirms expression of the EV component Alix. Scale bars: I,J = 5 μm.

The majority of EVs observed in the peripheral circulation and pericardial space are likely to be below the resolution limit of light-sheet or confocal microscopy (≥250 nm) meaning live imaging analysis cannot determine their true size. Additionally, the point spread function of the imaging system results in the particles appearing larger than they are. Nevertheless, the fluorescence confirms their cellular origin and allows them to be visualised even if their true size needs to be investigated via other methods. To investigate the endogenous EVs further, we analysed isolated fluorescently labelled EVs from Tg(*actb2:HRAS-EGFP*); Tg(*kdrl:mCherry-CAAX*) double transgenic larvae by high resolution TIRF microscopy (approx. max resolution <100 nm) together with 60 nm synthetic beads revealing *actb2*(GFP)+ EVs of a similar size to the synthetic beads (Figure 3I,J). Similar to the synthetic nanoparticles already described (Figure IIIE in the Data Supplement), these beads (and likely many EVs) are below the resolution limit of light microscopy, however, they can still be visualised by their fluorescence, although their true size still cannot be accurately determined by these methods. TIRF imaging of a single sorted (FAVS) vesicle confirms triple labelling for calcein (intact vesicle), GFP (near-ubiquitous marker) and mCherry (endothelial specific) (Figure 3J). Finally, western blot analysis with an antibody against known EV component ALIX, reveals expression in isolated small and large EV fractions (Figure 3K).

Collectively, our data describe methods to label cell-type specific EVs *in vivo* and highlights larval zebrafish as an ideal model to investigate EV function in multiple different cardiovascular functions and pathologies.

### Tissue injury models affect endogenous EVs *in vivo*

We next sought to demonstrate the utility of this labelling system to observe changes to endogenous vesicle production during pathologic states. As tissue hypoxia is an integral component of ischaemic injury and has been shown to induce release of EC-EVs primarily *in vitro* ^53^, we used our model to investigate this *in vivo*. Incubating 3 dpf Tg(*kdrl:mCherry-CAAX*) larvae in 5% oxygen for 18 hours significantly increased the number of EC-EVs observed in the peripheral circulation when compared to control larvae maintained in normoxic conditions (Figure VA-C and Video IX in the Data Supplement). This suggests that a global ischaemic/hypoxic environment increases EC-EV release into the peripheral circulation indicating dynamic EV responses following non-invasive tissue challenge in larval zebrafish. This further demonstrates the potential of the larval zebrafish model to assess EV function in pathological states *in vivo*. To further address the full complexity of endogenous cardiovascular EVs released following localised cardiac injury in fully differentiated tissues, we next investigated EVs in adult zebrafish before and after a well-established model of cardiac injury ^38^.

### Characterisation of EVs from adult cardiac tissue

To characterise EVs from adult cardiac tissue we extracted calcein+ and CM-EVs from isolated ventricles of uninjured adult Tg(*myl7:HRAS-mCherry*) fish and analysed these *ex vivo* using similar techniques described for larval samples (Figure 4A). To validate these adult cardiac EVs and confirm their true size, we performed TEM revealing EVs with a characteristic cup-like structure in sizes ranging from ~50-300 nm (Figure 4B-D), supporting our DLS data on size distribution of EVs in larvae. Flow cytometry analysis revealed populations of intact, cell-type specific CM-EVs derived from adult ventricles that could be obtained in much higher numbers than from entire larval fish (Figure 4E-I). NTA demonstrated calcein+ and CM-EVs with similar size ranges before and after FAVS with this analysis further confirming their lipid composition (Figure 4J-M). Finally, to determine if cell type specific EVs could be visualised *in vivo* in adults, we performed live imaging of superficial vessels of adult Tg(*kdrl:mCherry-CAAX*) fish revealing EC-EVs present in the peripheral circulation (Figure 4N and Video X in the Data Supplement).

**Figure 4 -.**
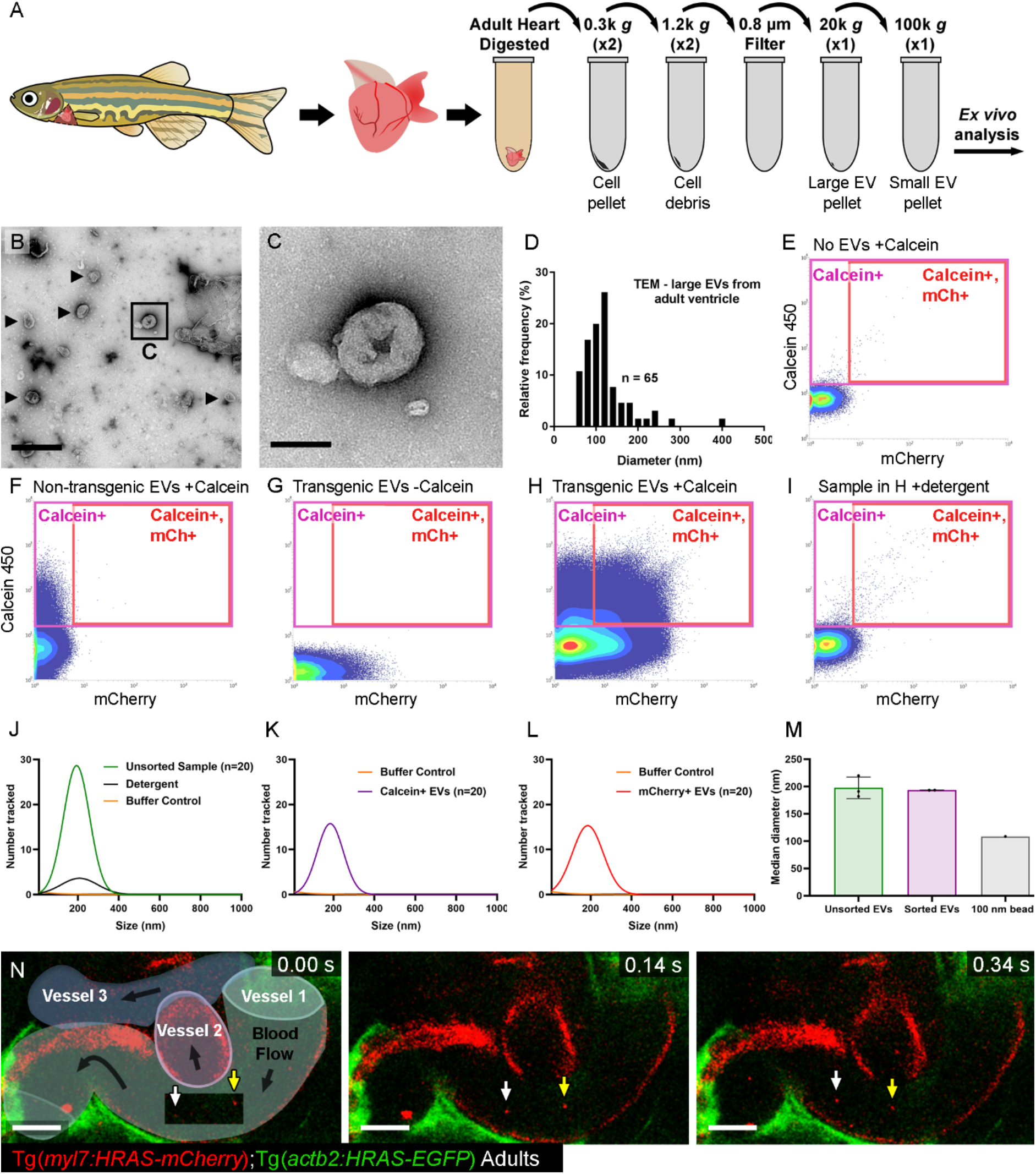
Validation of endogenous cardiovascular EVs from adult zebrafish. **(A)** Schematic of an adult zebrafish, adult heart and centrifugation process to obtain adult cardiac EV fractions. (**B,C**) TEM negative stain micrograph of an isolated large EV fraction from a pool of adult ventricles (n = 3). C shows a higher magnification view of the boxed region in B. (**D**) Histogram of the size distribution of EVs from a large EV pellet from adult zebrafish ventricles analysed by TEM. n = number of vesicles. (**E-I**) Typical flow cytometry scatter plots showing the gates used to sort adult cardiac EVs and the controls used to define these gates. (**J-L**) Gaussian distribution of NTA on an unsorted small EV fraction before and after detergent treatment (J), the same sample following sorting for calcein+ EVs (K) and the same sample following sorting for mCherry+ calcein+ EVs (L). (**M**) Median diameter measurements from NTA of unsorted and sorted EVs (data from J-L, sorted is K,L combined), including 100 nm reference beads. Unsorted and sorted EVs are of a similar size. (**N**) Schematic overlay describing the position of the three vessels visible in the integrated time series of live imaging of EC-EVs in the peripheral circulation of an adult Tg(*actb2:HRAS-EGFP*); Tg(*kdrl:mCherry-CAAX*) double transgenic fish. White and yellow arrows indicate two EC-EVs moving with the blood flow. Scale bars: B = 500 nm; C = 100 nm; N = 10 μm.

### Exchange of EVs differs between cardiac cell types

Next, we sought to investigate the transfer of endogenous EVs between different cell types within the cardiac microenvironment. We performed confocal imaging of hearts from combinations of transgenics allowing us to observe EV transfer between cardiomyocytes, ECs and macrophages (Figure 5). Combining membrane tethered mCherry lines (cells and EVs labelled) with cytoplasmic GFP lines (only cells labelled) revealed transfer of CM-EVs to ECs (Figure 5A-D), but indicated only very limited transfer of EC-EVs to cardiomyocytes (Figure 5E-H). Conversely, transfer of both CM-EVs and EC-EVs to macrophages resident in the heart was observed (Figure 5I-P and Video XI in the Data Supplement). Quantification of the degree of transfer between different cell types suggests the majority of this labelled material is received by macrophages, followed by ECs (Figure 5Q,R), with the lowest transfer observed to cardiomyocytes (Figure 5R). These data reveal new insights into cellular communication occurring *in vivo* between different cardiovascular cell types and demonstrate the utility of the labelling method to investigate complex cellular processes in different tissues.

**Figure 5.**
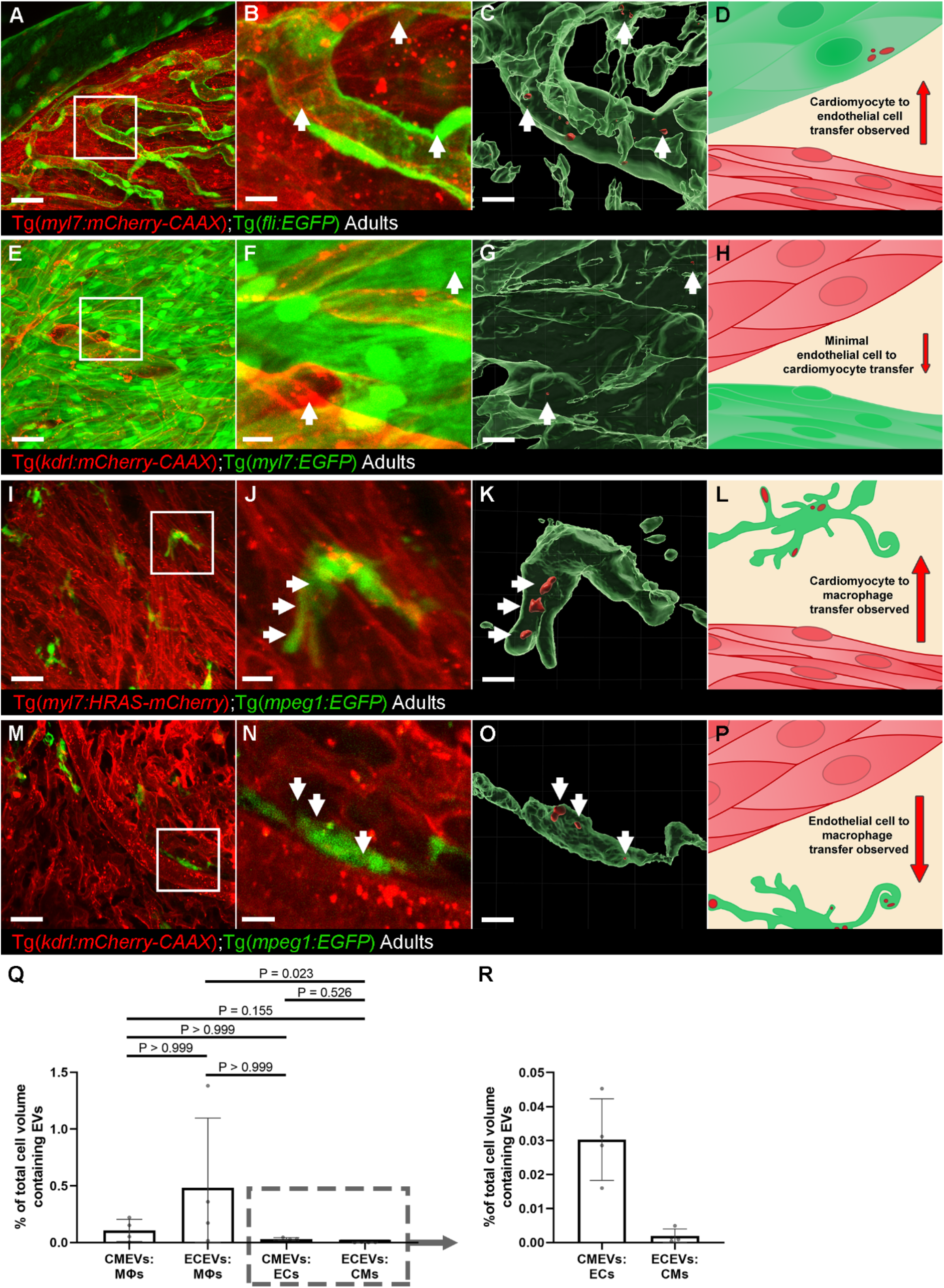
Whole-mount high resolution imaging of fixed adult hearts to characterise EV transfer to recipient cells. (**A,E,I,M**) Overview maximum projection images showing the surface of adult Tg(*myl7:HRAS-mCherry*); Tg(*fli:EGFP*) (A), Tg(*kdrl:mCherry-CAAX*); Tg(*myl7:EGFP*) (E), Tg(*myl7:HRAS-mCherry*); Tg(*mpeg1:EGFP*) (I) and Tg(*kdrl:mCherry-CAAX*); Tg(*mpeg1:EGFP*) (M) zebrafish ventricles. (**B,F,J,N**) Maximum projections of digital zooms of the boxed regions in A,E,I,M. (**C,G,K,O**) 3-D reconstruction of the images in B,F,J,N (membrane tethered fluorophore+ cells have been removed for clarity). (**D,H,L,P**) Schematic summaries for each row of potential EV transfer between the labelled cell types. (**Q,R**) Volume measurements of EV containing compartments within recipient cell types (lower x axis label), shown as a percentage of the total volume measurement for that cell type within a field of view. Taken from 3D image analysis. The boxed region in Q highlights the expanded view in R. Arrows indicate EVs within recipient cells. Scale bars: A,E,I,M = 20 μm; B,C,F,G,J,K,N,O = 5 μm.

### Cardiovascular EVs exhibit dynamic responses to cardiac injury

As EVs are linked to the progression of cardiovascular disease after myocardial injury ^54^, we next assessed the EV response to tissue damage and further demonstrate the value of the adult model. We performed cardiac cryoinjury on adult zebrafish and extracted EVs from isolated ventricles of unwounded and 24 hours post injury (hpi) fish and analysed them via flow cytometry and DLS (Figure 6). Flow cytometry did not reveal significant changes to the overall number of calcein+ or intact CM-EVs in the heart following cardiac injury (Figure 6A,B). By contrast, there were significantly fewer EC-EVs as a proportion of calcein+ EVs (Figure 6C), suggesting an increase in EVs derived from additional cell types e.g. interstitial fibroblasts or inflammatory cells, which were not assessed here. Interestingly, DLS revealed a significant shift in the size distribution of total EVs at 24 hpi when compared to unwounded hearts (Figure 6D) suggesting a dynamic EV response following cardiac injury.

**Figure 6.**
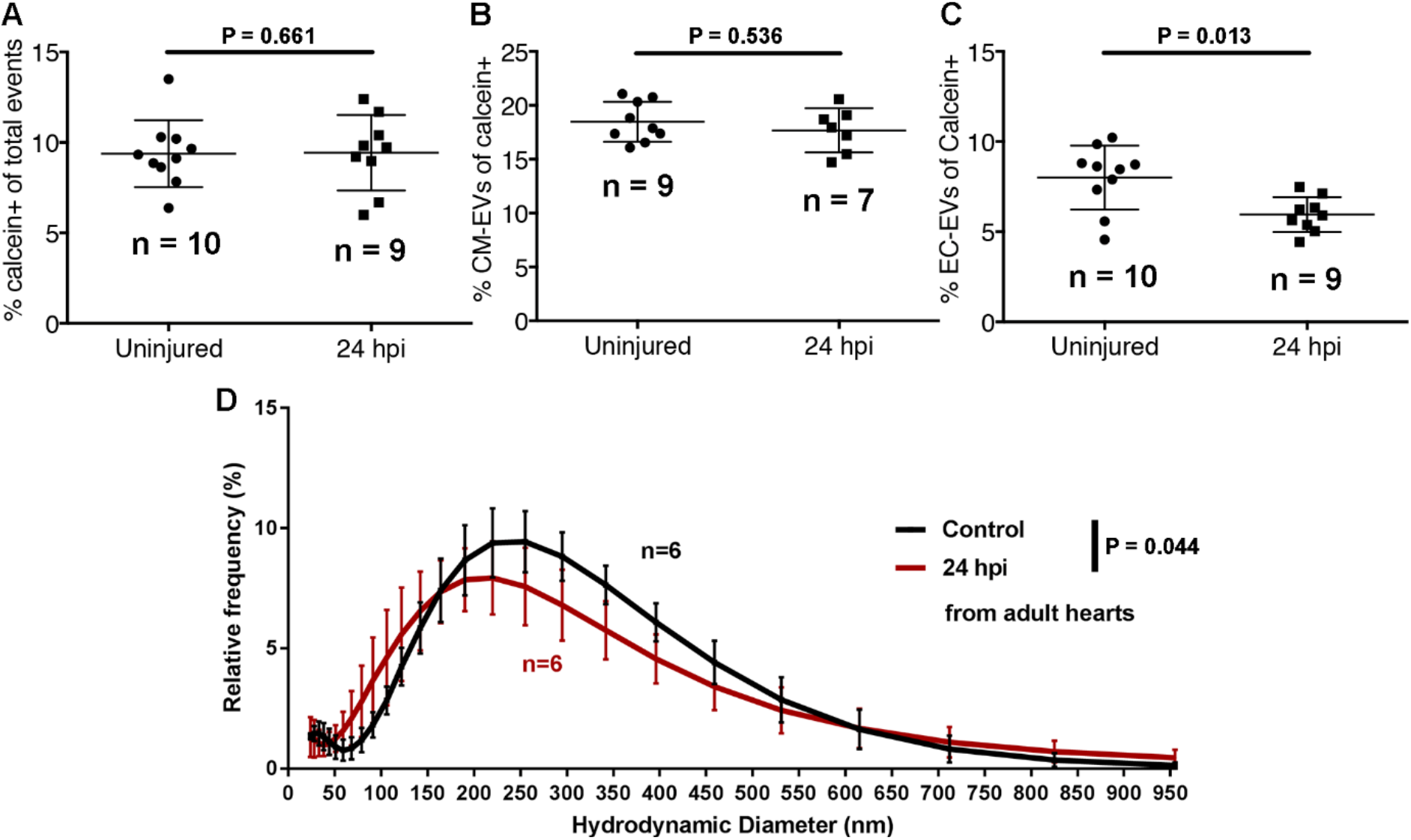
A model of cardiac injury induces dynamic changes in cardiovascular EVs from adult zebrafish. **(A-C)** Quantification of the number of calcein+ (A), *myl7*(mCherry)+ (B) and *kdrl*(mCherry)+ EVs in uninjured and injured hearts at 24 hpi. (**D**) Histogram of DLS analysis reveals a significant shift in the size of overall EVs at 24 hpi compared to uninjured hearts. Statistical analysis in A-C: Two-tailed Mann-Whitney tests. Statistical analysis in D: a custom permutation test using total variation distance was used to test the null hypothesis that control and 24 hpi distributions were the same.

## Discussion

Cell-cell communication roles for EVs have been mostly studied in *ex vivo* and *in vitro* cell models. However, relatively little is known about the *in vivo* function of native EVs. Here, we have described methods to fluorescently label cell-type specific endogenous EVs and visualised these *in vivo* in a vertebrate zebrafish model system. Our findings complement recent studies that describe the labelling, biodistribution and potential functional roles of exogenous tumour derived and endogenous CD63+ EVs in larval zebrafish ^25, 29^. The described techniques and advancements in the use of zebrafish for *in vivo* EV research pave the way for investigations into multiple homeostatic and pathophysiological states ^55^ and the involvement of endogenous EVs in cellular processes such as tissue regeneration.

The labelling strategy described here allows the identification of the cellular origin of EVs and their visualisation *in vivo* without reliance on the expression of specific marker proteins. Currently, the proportion of EVs that incorporate fluorophore labelling during biogenesis is incompletely understood, potentially limiting the number of EVs produced by each cell type that we can successfully identify ^56^. Indeed, flow cytometry assessments suggest that approximately 30-65% of events are labelled with calcein (e.g. intact vesicles that contain esterases) and that approximately 30% of these are labelled with *actb2*(GFP)+ (thought to be a near-ubiquitous promoter), suggesting relatively restricted fluorophore and esterase incorporation and/or limited preservation of intact EVs during isolation and/or sorting. Additionally, the relative incorporation of fluorophore between different EV subpopulations is currently not well determined and requires additional characterisation. Regardless of subgroup, shared fluorescence between the selected cell types and EVs observed within extracellular space confirm the cellular origin of these vesicles and sufficient numbers of intact vesicles can be obtained for further studies. Combining this system with other labelling methods will allow cell type specific sub classes of EVs to be labelled *in vivo* in the future.

We have also shown for the first time that EVs from adult zebrafish can be visualised and extracted. Using combinations of stable transgenic lines we can observe the conversations between different cell types in the cardiac microenvironment. Tissue-resident macrophages received the majority of labelled material from other cell types although ECs also received CM-EVs. Similarly, in larvae EC-EVs were observed within intravascular macrophages, which were often observed making protrusions, potentially to catch passing EVs. This suggests macrophages may receive the majority of EVs during homeostasis *in vivo*, supporting previous findings ^25, 29^. EC-EVs could be observed moving rapidly with the blood flow in superficial vessels of adult fish and TEM reveals a characteristic cup-shaped appearance of extracted vesicles from isolated adult hearts. TEM measurements and DLS analysis reveal similar size distributions for total EVs extracted from adult hearts and from whole larvae, and are in line with previous reports of EVs from zebrafish, mice and humans ^4, 25, 50, 57, 58^.

Importantly, we have demonstrated that models of cardiovascular injury can induce changes to EV number and size, suggesting a dynamic intercellular communication response that involves EVs, supporting previous *in vitro* models, clinical findings and studies in non-regenerative rodent models of MI ^4, 5, 10, 57, 59–61^. Similar numbers of overall EVs and CM-EVs following cardiac injury but a reduction in EC-EVs, as a proportion of total (calcein+) EVs, suggest an increased proportion of EVs deriving from an, as yet undetermined, cell-type. Interestingly, we observed an increase in EC-EVs following larval injury models and a reduction in EC-EVs following an adult MI model, suggesting dynamic changes to EVs that may be highly tissue- and time-point dependent. Further studies will be required to determine if the reduction in EC-EVs results from reduced numbers of ECs in the injured heart at 24 hpi or from dynamic changes to EC-EV production following injury. Our data further suggest that there is a significant shift to smaller EVs at 24 hpi potentially indicating increased exosome release. This is similar to what has been described for clinical plasma samples in patients undergoing cardiac surgery ^4^ and may represent an active shift in EV biogenesis in response to MI. Additionally, further studies to define the cargo and composition of these post-ischaemic EVs could reveal their roles in tissue repair and regeneration and identify potential therapeutic targets.

In summary, we have shown that zebrafish are an ideal model to investigate endogenously produced cell-type specific EVs stably labelled with fluorophores, revealing their cellular origin and allowing them to be observed and tracked *in vivo*. Large numbers of cell-type specific EVs can be obtained from adult hearts allowing future evaluations of cargo during homeostasis and in models of disease. Finally, multiple cell types exchange EVs in the adult heart and models of ischaemic injury produce dynamic changes to EV size and numbers produced by different cell types demonstrating the role of these small vesicles in these processes.

## Supporting information

Supplemental Video III

Supplemental Video VII

Supplemental Video VIII

## Acknowledgements

The authors wish to thank the Wolfson Bioimaging Facility for imaging expertise, Professor Dan Peer (Tel Aviv University) for providing synthetic nanoparticles, and Professor Jonathan Rougier (University of Bristol) for statistical advice.

## Abbreviations

EV: Extracellular vesicle
EC: Endothelial cell
dpf: Days post fertilization
DLS: Dynamic light scattering
NTA: Nanoparticle tracking analysis
FAVS: Fluorescence activated vesicle sorting
TEM: Transmission electron microscopy
TIRF: Total internal reflection fluorescence
EC-EV: Endothelial cell-derived EV
DA: Dorsal aorta
CHT: Caudal haematopoietic tissue
CM-EV: Cardiomyocyte-derived EV
hpi: Hours post injury

## Highlights

- Multiple cell types produce EVs in larval and adult zebrafish *in vivo*, including endothelial cells and cardiomyocytes
- A transgenic membrane-tethered fluorophore system allows EVs from specific cell types to be visualised, tracked and obtained for downstream investigations.
- EVs are transferred between multiple different cell types in the adult zebrafish heart,
- The production of endogenous EVs is altered following tissue injury.

## Supplemental Figures

**Supplemental Figure I.**
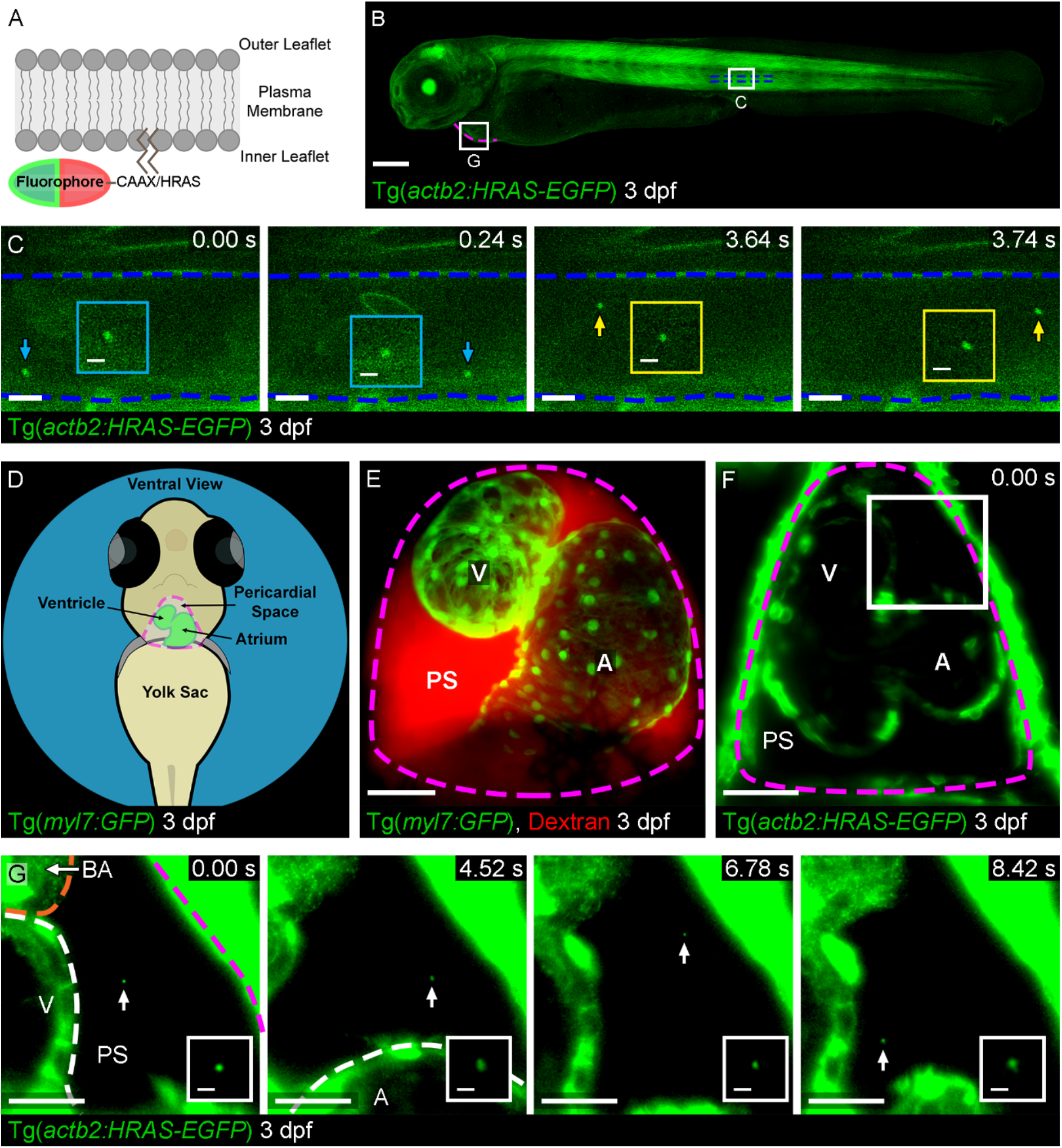
EV labelling strategy and live imaging in the peripheral circulation and pericardial space. (**A**) Schematic representation of the cell/EV labelling strategy. The fluorophore is tethered to the inner leaflet of the plasma membrane via a CAAX or HRAS motif. (**B**) Overview image of a Tg(*actb2:HRAS-EGFP*) larval zebrafish at 3 dpf. The boxed areas define the regions shown in the image sequences in C and G, as indicated. (**C**) Image sequence of GFP labelled EVs (arrowed) moving through the DA. Blue dashed lines demark the endothelium lining the vessel. Insets show higher magnification views of the arrowed EV. (**D**) Schematic showing the position of the larval heart and pericardial space in a ventral view of a 3 dpf fish. (**E**) Image of the heart in ventral view with fluorescently labelled cardiomyocytes of a Tg(*myl7:GFP*) fish (green) and injected Dextran to demonstrate the extent of the pericardial space (red) at 3 dpf. (**F**) Overview image of the heart of a Tg(*actb2:HRAS-EGFP*) larval zebrafish at 3 dpf. (**G**) Image sequence of the boxed region in F showing a GFP labelled EV (arrowed and inset) moving through the pericardial space. The magenta dashed line in B,D-G demarks the outer pericardial wall. The orange dashed line in G demarks the bulbus arteriosus. The white dashed line in G demarks the ventricle and atrium. Anterior is to the left in B,C. BA = bulbus arteriosus, V = ventricle, A = Atrium, PS = Pericardial space. Scale bars: B = 200 μm; C = 5 μm; insets in C,G = 2 μm; E,F = 50 μm; G = 20 μm.

**Supplemental Figure II.**
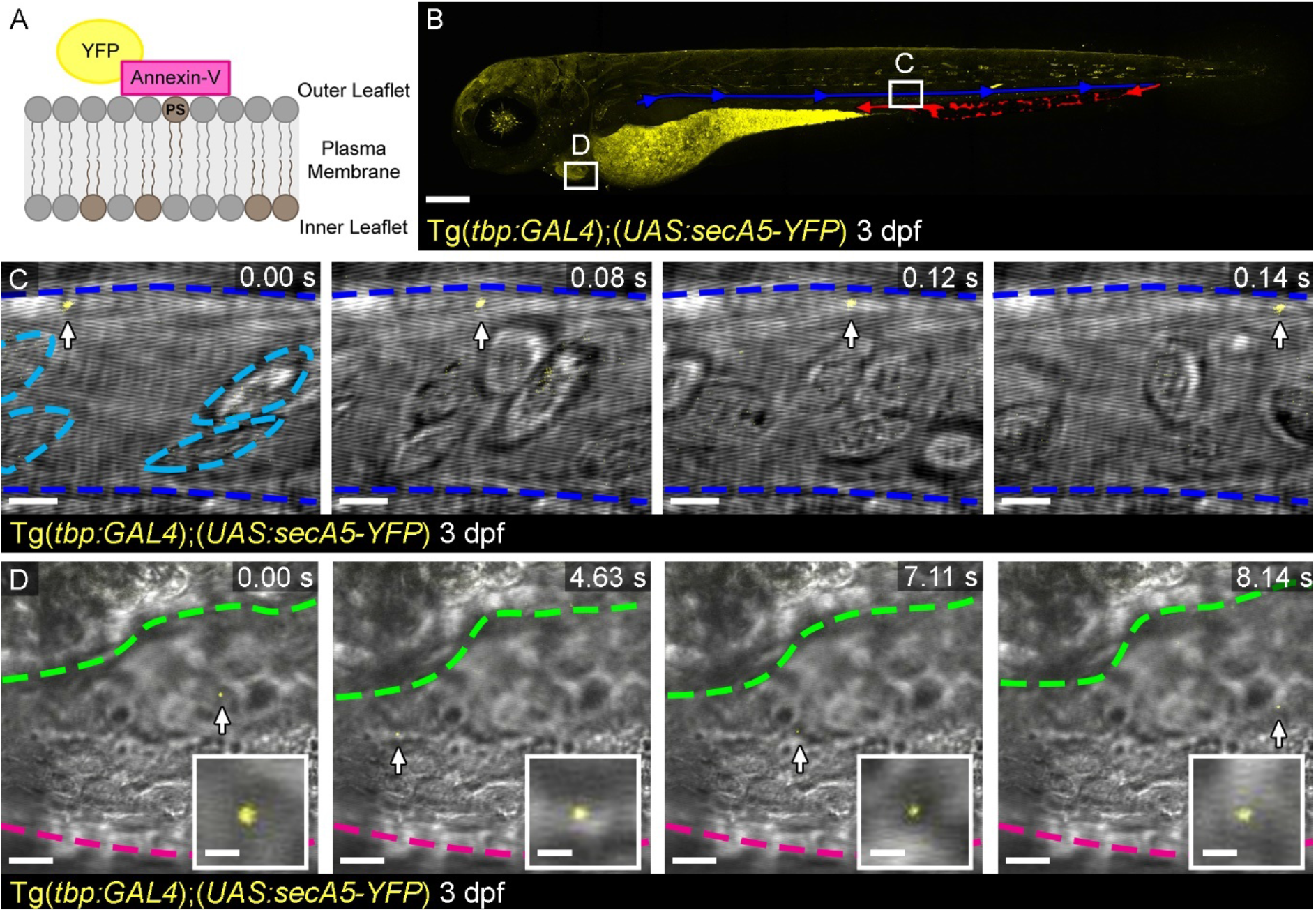
Live imaging of a global EV labelling strategy via secreted Annexin-V. (**A**) Schematic representation of the Annexin-V labelling strategy. (**B**) Overview image of a larval Tg(*tbp:GAL4*);(*UAS:secA5-YFP*) zebrafish at 3 dpf. Blue indicates the DA and red the caudal haematopoietic (venous) tissue (CHT). Boxes depict the approximate position of the image sequences in C and D. (**C**) Image sequence of an Annexin-V labelled EV moving through the DA. Blood cells can be clearly seen in brightfield. (**D**) Image sequence of an Annexin-V labelled EV (also inset) moving through the pericardial space. The light blue dashed lines in C demark blood cells. The magenta dashed line in D demarks the outer wall of the pericardial space, green outlines the ventricle. Anterior is to the left. Scale bars: B = 200 μm; C = 5 μm; D = 10 μm; insets in D = 2 μm.

**Supplemental Figure III.**
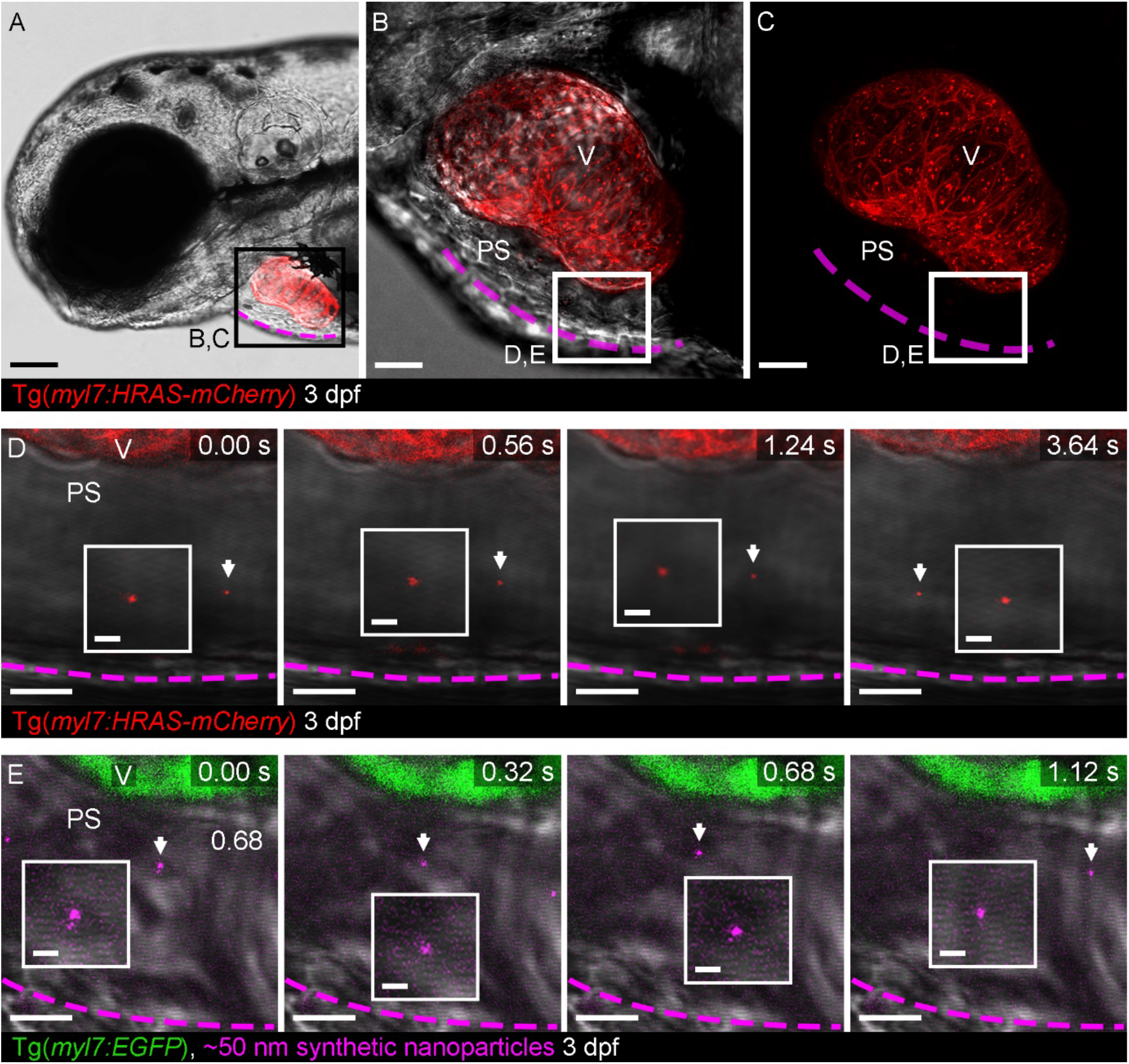
Live imaging cell-type specific EVs and 100 nm synthetic nanoparticles in the pericardial space. (**A**) Lateral anterior view of a Tg(*myl7:HRAS-mCherry*) larval zebrafish at 3 dpf. The boxed area defines the approximate region shown in B,C. (**B,C**) Optical zoom of the heart shows the entire ventricle; with brightfield (B) and without (C). Boxed area defines the approximate regions shown in D,E. (**D**) Image sequence of higher magnification views of the ventral pericardial space. mCherry+ CM-EVs are observed moving freely through the pericardial space at a distance from the pericardial wall (arrowed and inset). (**E**) Image sequence of higher magnification views of the ventral pericardial space of a Tg(*myl7:GFP*) larvae at 3 dpf. Cy5+ 100 nm synthetic nanoparticles are observed moving freely through the pericardial space as observed for CM-EVs (arrowed and inset). The magenta dashed line in A-E demarks the outer pericardial wall. Anterior is to the left. V = ventricle, PS = Pericardial space. Scale bars: A = 100 μm; B,C = 25 μm; D,E = 10 μm; insets in D,E = 2 μm.

**Supplemental Figure IV.**
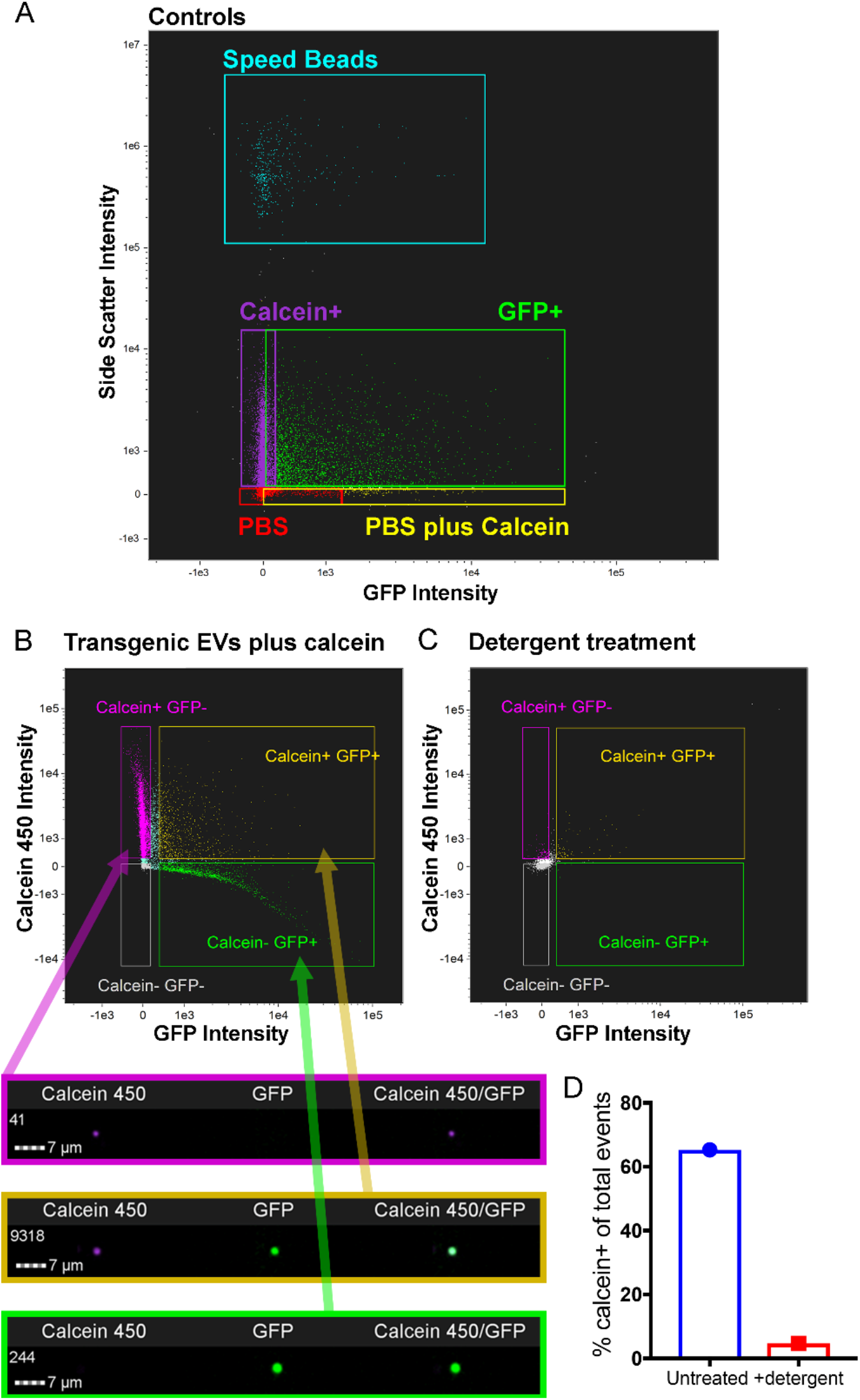
Fluorescent gating strategy and ImageStream analysis of larval zebrafish EVs. (**A-D**) ImageStream analysis and quantification of EVs from a pool of 30 whole Tg(*actb2:HRAS-EGFP*) larvae. Control experiments using PBS, PBS plus calcein AM, *actb2*(GFP+) EVs with and without calcein AM and speed beads (1.5 μm diameter, carboxylated polystyrene microspheres, see methods for details) were used to set correct gates for double positive (fluorescence and calcein) EVs (A). Analysis of EVs from Tg(*actb2:HRAS-EGFP*) fish demonstrates different populations of EVs which can be individually visualised (lower panels) (B). Detergent treatment destroys EV integrity further confirming their lipid nature (C,D).

**Supplemental Figure V.**
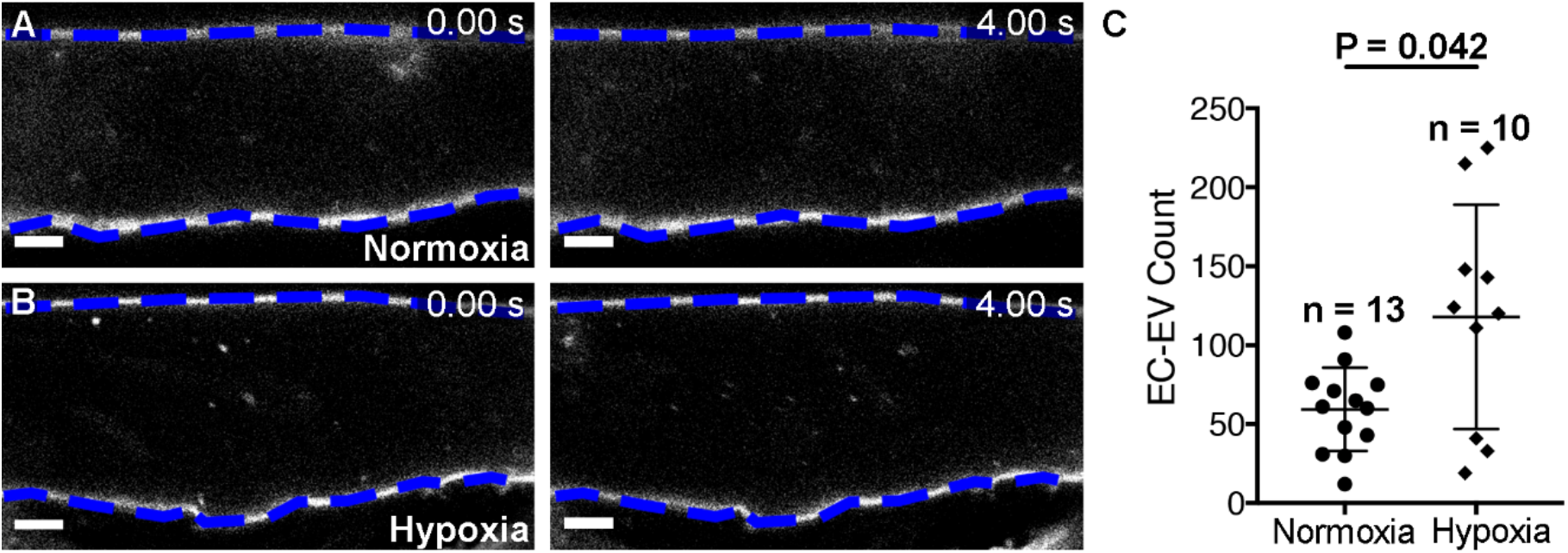
EV response to hypoxia in larval zebrafish. (**A,B**) Image sequences of the DA of a 4 dpf Tg(*kdrl:mCherry-CAAX*) fish under normoxic (A) or hypoxic (5% oxygen) conditions (B). (**C**) Quantification of the number of EC-EVs in the DA under normoxic or hypoxic conditions. Statistical analysis in C: Two-tailed Mann-Whitney test. Scale bars: A,B = 5 μm.

## Supplemental Videos

**Supplemental Video I. Live imaging of *actb2+* EVs in the peripheral circulation of a larval zebrafish.** Image sequence (751 frames = 15 seconds) of *actb2+* EVs passing through the DA of a 3 dpf Tg(*actb2:HRAS-EGFP*) larval zebrafish. Scale bar = 5 μm.

**Supplemental Video II. Live imaging of *actb2+* EVs in the pericardial space of a larval zebrafish.** Image sequence (191 frames = 3.28 seconds) of *actb2+* EVs travelling through the pericardial fluid of a 3 dpf Tg(*actb2:HRAS-EGFP*) larval zebrafish. Scale bar = 50 μm, 5 μm.

**Supplemental Video III. Live imaging of Annexin-V labelled EVs in the pericardial space and peripheral circulation of larval zebrafish.** Image sequence (13 frames = 0.26 seconds) of *YFP+* EV travelling through the DA of a 3 dpf Tg(*tbp:GAL4*);(*UAS:secA5-YFP*) larval zebrafish. Image sequence (327 frames = 13.08 seconds) of *YFP+* EVs travelling through the pericardial fluid of a 3 dpf Tg(*tbp:GAL4*);(*UAS:secA5-YFP*) zebrafish. Scale bar = 5 μm, 10 μm.

**Supplemental Video IV. Live imaging of EC-EVs in the peripheral circulation of a larval zebrafish.** Image sequence of mCherry+ EC-EVs passing through the DA (751 frames = 15 seconds) and the CHT (751 frames = 15 seconds) of a 3 dpf Tg(*kdrl:mCherry-CAAX*) larval zebrafish. Scale bar = 5 μm.

**Supplemental Video V. Live imaging of an intravascular macrophage in the peripheral circulation of a larval zebrafish.** Image sequence (451 frames = 9 seconds) of a GFP+ macrophage containing mCherry+ EC-EVs in the CHT of a 3 dpf Tg(*kdrl:mCherry-CAAX*); Tg(*mpeg1:EGFP*) larval zebrafish. Note the protrusion into the peripheral circulation. Scale bar = 10 μm.

**Supplemental Video VI. Live imaging of CM-EVs in the pericardial space of a larval zebrafish.** Image sequence (669 frames = 12 seconds) of mCherry+ CM-EVs travelling through the pericardial fluid of a 3 dpf Tg(*myl7:HRAS-mCherry*) larval zebrafish. Scale bar = 50 μm, 5 μm.

**Supplemental Video VII. Z-stack of the pericardial space of a Tg(*myl7:GFP*) larval zebrafish injected with dextran.** Slice-by-slice view (interval = 0.99 μm, total z-depth = 167.31 μm) of the pericardial space of a Tg(*myl7:GFP*) 3 dpf larvae injected with dextran. Limited cells are seen within the pericardial space. Scale bar = 50 μm.

**Supplemental Video VIII. Live imaging CM-EVs and 100 nm synthetic nanoparticles in the pericardial space of a larval zebrafish.** Image sequence of mCherry+ CM-EV (260 frames = 5.2 seconds) and 100 nm synthetic nanoparticles (48 frames = 12.96 seconds) travelling through the pericardial space of a 3 dpf Tg(*myl7:HRAS-mCherry*) and Tg(*myl7:GFP*) larval zebrafish respectively. Scale bar = 10 μm.

**Supplemental Video IX. Live imaging sequence of EC-EVs in the peripheral circulation of larval zebrafish hypoxia-induced injury models.** Image sequences (522 frames = 10.44 seconds) of EC-EVs passing through the DA of 4 dpf Tg(*kdrl:mCherry-CAAX*) larval zebrafish under normoxic and hypoxic conditions (18 hours at 5% Oxygen). Scale bar = 5 μm.

**Supplemental Video X. Live imaging sequence of EC-EVs in the peripheral circulation of adult zebrafish.** Image sequences (837 frames = 16.62 seconds) of mCherry+ EC-EVs passing through a superficial blood vessel near the gills of a 12-month Tg(*actb2:HRAS-EGFP*); Tg(*kdrl:mCherry-CAAX*) adult zebrafish. Scale bar = 5 μm.

**Supplemental Video XI. 3D reconstruction and rendering of the ventricular tissue of a Tg(*kdrl:mCherry-CAAX*); Tg(*mpeg1:EGFP*) adult zebrafish.** 3D view of mCherry+ EC-EVs within cardiac macrophages of a Tg(*kdrl:mCherry-CAAX*); Tg(*mpeg1:EGFP*) adult zebrafish heart. Images were reconstructed and rendered using Imaris software. Scale bar = variable with zoom, see video.

